# Distinct seasonal acclimatisation trajectories characterize transplanted and natural meadow seagrass plants

**DOI:** 10.64898/2026.08.14.744801

**Authors:** Gianmarco Valenti, Alberto Sutera, Francesco Cosenza, Fabio Badalamenti, Vincenzo Maximiliano Giacalone, Francesco Carimi, Francesco Mercati, Guglielmo Puccio, Roberto De Michele

## Abstract

Successful establishment is a critical determinant of seagrass restoration, yet the molecular mechanisms underlying seedling acclimatisation to natural environments remain poorly understood. Here, we combined seasonal physiological observations, transcriptome profiling, and gene co-expression network analysis to investigate the mechanisms underlying the early post-transplantation phase of *Posidonia oceanica*, a dominant foundation seagrass species, following transplantation. Transplanted seedlings were compared with plants from adjacent natural meadows over the first six months after transplantation using leaf and root samples collected in spring, summer, and autumn. Tissue identity was the primary driver of transcriptomic variation, but transplanted seedlings remained transcriptionally distinct from plants in natural meadows throughout the study, with roots showing greater divergence than leaves, suggesting tissue-specific trajectories of post-transplantation acclimatisation. The early post-transplantation phase was characterised by the activation of genes associated with RNA processing, transcriptional regulation, and abscisic acid signalling. During a summer marine heatwave (28 °C), both plant groups induced conserved heat-response pathways, including heat-shock proteins and protein-folding mechanisms. Furthermore, transplanted seedlings maintained higher expression of genes involved in photosystem II repair and photoprotection and exhibited reduced leaf growth and extensive leaf necrosis, consistent with a greater requirement for photosynthetic maintenace under prolonged thermal stress. Gene co-expression network analysis revealed that regulatory networks governing structural integrity, hormone signalling, and defence were more stable in natural meadow plants, while transplanted seedlings progressively reorganized their gene co-expression patterns to resemble those of natural meadow plants, particularly in leaves. Our findings reveal tissue-specific molecular trajectories of acclimatisation during early seedling establishment and identify candidate molecular indicators of field acclimatisation and thermal stress responses, providing new mechanistic insights relevant to seedling-based seagrass restoration under climate change.

## 1. Introduction

*Posidonia oceanica* (L.) Delile is the dominant foundation seagrass of the Mediterranean Sea, where it forms extensive meadows from the shoreline to depths of approximately 40 m (Telesca et al. 2015). These meadows rank among the most productive and ecologically valuable coastal ecosystems in the region, sustaining marine biodiversity, enhancing ecosystem functioning and contributing to the maintenance of coastal environmental stability (Scanu et al. 2022). *P. oceanica* is a monoecious species that propagates both vegetatively and sexually through flowering, fruit production, seed dispersal, anchorage, stabilization and germination (Guerrero-Meseguer et al. 2018). Despite its ecological importance, *P. oceanica* meadows have faced a dramatic regression over recent decades, with an estimated decline of approximately 34%, primarily driven by increasing anthropogenic pressures and climate change (Telesca et al. 2015). Consequently, effective conservation and restoration strategies are urgently needed to preserve the ecological functions and ecosystem services provided by this foundation species.

Current restoration programs are predominantly based on the transplantation of vegetative cuttings into degraded meadows (Alagna et al. 2019; Calvo et al. 2021; Bacci et al. 2025). Although effective under certain conditions, this approach presents several limitations, including damage to donor meadows, reduced genetic diversity and high economic and logistics costs (Seddon 2004). Seed-based restoration has recently emerged as a promising alternative because it exploits the abundant availability of beach-cast fruits and seeds while preserving donor meadows and promoting greater genetic diversity within restored populations (Escandell-Westcott et al. 2023). Importantly, stranded fruits retain viable seeds, facilitating the collection of propagules suitable for restoration (Sutera et al. 2024), and recent methodological advances have established protocols for the long-term storage of *P. oceanica* seeds, overcoming the seasonal limitation in propagule availability (Sutera et al. 2025). Regardless of the restoration strategy employed, however, successful establishment ultimately depends on the ability of transplanted plants or seedlings to rapidly acclimate to their new environment through coordinated molecular and physiological responses.

Environmental acclimatisation requires extensive transcriptional reprogramming that enables plants to perceive external cues, adjust their metabolism and maintain cellular homeostasis under changing conditions. In seagrasses, this process is expected to be highly tissue specific. Leaves are primarily responsible for photosynthetic carbon assimilation and the perception of fluctuations in light and water-column conditions, whereas roots directly interact with the sediment, where they mediate anchorage, nutrient acquisition and belowground environmental sensing (Valenti et al. 2026). Understanding how these functionally distinct tissues coordinate their transcriptional responses during the early post-transplantation phase is therefore essential for identifying the molecular processes associated with field acclimatisation and potentially relevant to restoration outcomes.

The recent availability of the *P. oceanica* genome, together with those of other seagrass species, has substantially expanded our understanding of the genomic innovations associated with the evolutionary transition of angiosperms from terrestrial to marine environments (Ma et al. 2024). However, genomic resources alone provide only limited insight into the dynamic regulatory processes that enable plants to respond to environmental change. To date, transcriptomic studies in *P. oceanica* have largely focused on adult plants exposed to specific abiotic stressors under controlled laboratory conditions (Marín-Guirao et al. 2019; Ruocco et al. 2021; Pazzaglia et al. 2022; Santillán-Sarmiento et al. 2023). Even studies conducted under natural field conditions have exclusively investigated established meadows (Dattolo et al. 2013; Entrambasaguas et al. 2017; Procaccini et al. 2017). In contrast, only two studies have examined transcriptomic regulation in seedlings, both under controlled mesocosm conditions: our recent characterization of developmental transcriptomic dynamics (Valenti et al. 2026) and the transcriptional responses of seedlings to warming and ocean acidification (Pazzaglia et al. 2025). Consequently, the molecular mechanisms governing the transition from controlled cultivation to natural environmental conditions remain largely unexplored.

Here, we address this knowledge gap by providing the first transcriptomic characterization of *P. oceanica* seedlings during the first six month following transplantation into a natural environment. By comparing tissue-specific transcriptomes of leaves and roots from transplanted seedlings and neighbouring mature meadows across successive seasons, we investigated how transcriptional programs changed during early field acclimatisation. We specifically asked whether the expression profile of transplanted seedlings became progressively more similar to those of natural plants, which biological processes and co-expression modules were associated with temporal changes, and how seasonal environmental variation was reflected in tissue-specific transcriptional responses. This work provides a molecular framework for understanding post-transplantation acclimatisation and offers new insights relevant to the development of seedling-based restoration strategies for Mediterranean seagrass ecosystems.

## 2. Materials and methods

### 2.1 Seedling growth and experimental design

Seeds were collected from stranded fruits along the shore of Trabia, Sicily (37.994 N, 13.666 E) on May 12, 2021. The fruit husk was removed and the seeds cleaned from debris directly at the beach. Seeds were transported to the laboratory in refrigerated mobile tanks with seawater. In the laboratory, they were rinsed and stored at 4°C in artificial seawater (ASW, 3.8% salinity). Two days later, about 200 seeds were transferred to an aquarium containing 40 L refrigerated ASW maintained at 18 °C, equipped with biological filtration and growth light (300 lx with a photoperiod of 14 h/10 h light/dark), and no substratum. Seedlings were grown in these conditions for almost a year (Figure S1A). Dead material was removed as needed, and ASW was replaced weekly during the first two months, and then monthly. On April 26, 2022, the seedlings were placed in 5 cm diameter pots filled with pebble stones (1-3 cm diameter) and secured with elastic net to prevent displacement (Figure S1B) (Zenone et al. 2025). Pots were attached in pairs through plastic bands and returned in the aquaria in upright position. The day after, pots were transported in refrigerated mobile tanks filled with ASW to the transplantation sites, located along the coast of Castellammare del Golfo, Sicily (site 1: 38°02’07.7”N 12°52’29.1”E; site 2: 38°02’45.4”N 12°52’14.1”E, Figure S2), using a diving boat. The sites were about 20-50 m from the shore, along an east-facing rocky outpost, at 5 m depth. Two pot pairs (total four pots) were secured with plastic bands in the upper section of each 50 cm long rebar. The rebars were driven deeply into the seabed, with the pots positioned at seabed level and partially embedded in the substrate by a few centimetres, adjacent to existing *P. oceanica* meadows (Figure S1C-E, Video S1) (Domínguez et al. 2012). Two groups of three rebars were transplanted in each site, signaled by buoys to ease their localization at subsequent collection times. Groups were 10 m distant. In total, 24 pots were transplanted in each site. The two sites were about 1 km apart and presented similar features. The first sampling (T1) was conducted one month later (May 26, 2022), with the collection of 2 rebars per site (one per group), corresponding to 16 seedlings total (Figure S1F). In the boat, seedlings were rapidly removed from the pot and photographed. Epiphytes, when present, were gently scraped off leaves. For transcriptomic analyses, we preserved tissues from only three seedlings per rebar, out of four. About 10 cm segments of a single root and leaf (the second innermost) were excised and preserved, separately, in 15 mL tubes with 10 mL of RNAlater (ThermoFisher). The second sampling (T2), three months after planting (July 26, 2022); and the third and final sampling (T3), six months after planting (October 25, 2022), followed the same procedure. Additionally, during the entire experimental period, 3 bundles per site at every time point were collected from existing, mature meadows for transcriptomic analysis of adult, established plants. For T2, samples from natural meadows in site 1 were accidentally destroyed, therefore we processed only the three replicates from site 2. From the bundles, 10 cm segments from roots and leaves were preserved in RNAlater as described above. Then, samples were stored at −80°C until further processing. In total, we collected 72 samples from seedlings and 30 from natural meadows (Table S1).

### 2.2 Physiological measurements and statistics

The number of leaves and roots was calculated from photographs of the 16 seedlings collected at each time point, regardless of their health status, before tissue excision. Four additional seedlings at the transplantation time (T0) were equally photographed and analysed, and then discarded. Maximum and total leaf and root lengths were measured with ImageJ (version 1.54). At each time point, all data were pooled, regardless of site and group of transplantation. Statistical significance was evaluated by Kruskal-Wallis test, followed by pairwise Mann-Whitney test (alpha = 0.05).

### 2.3 Environmental data collection

At transplantation and at each sampling time point, water samples were collected in proximity to the seedlings (Figure S1G). Using sterile plastic bottles, 1 L seawater was collected for chemical measurements and 0.5 L for microbiological tests at each site. The analyses were performed by Alimenti e Ambiente S.r.l. (Bagheria, Italy) on the same day of collection.

Daily mean air temperature was retrieved by the astronomical observatory of the University of Palermo (http://meteo.oapa.inaf.it/public/) (Figure S3). Sea water temperature, at the transplantation site at depth of 5.46 m, was estimated by E.U. Copernicus Marine Service (https://doi.org/10.48670/mds-00359) (Figure S3).

### 2.4 RNA extraction and analysis

Leaf and root samples were thawed, rinsed and blotted on filter paper, then ground in a mortar in liquid nitrogen. The ground material was processed using the Plant Tissue section of the Aurum™ Total RNA Mini Kit (BIO-RAD, Hercules, CA, USA), following the manufacturer’s protocol. Total RNA was quantified by g a Synergy Plate Reader spectrophotometer (BioTek), and its integrity checked by gel electrophoresis.

Library sequencing was performed on an Illumina NovaSeq 6000 platform, producing high-quality paired-end reads (Biodiversa, Italy). The initial quality assessment of the raw data was conducted using FastQC(Wingett and Andrews 2018). To ensure data integrity, raw reads were processed with TRIMMOMATIC to remove adapter sequences and low-quality bases (Sewe et al. 2022). This preprocessing step yielded a refined dataset of clean reads characterized by a median Phred quality score of 36. Sequencing revealed a skewed GC content, indicating the presence of eubacterial DNA. This was expected, as plants were covered by biofilm and *P. oceanica* seeds are known to harbour a diverse community of endophytes (Crucitti et al. 2026). Accordingly, reads were mapped to the *P. oceanica* reference genome (Ma et al. 2024) to exclude non-*P. oceanica* sequences from subsequent analyses. Mapping was executed using the STAR (Dobin et al. 2013) aligner with default settings. Finally, the quantification of gene expression was performed by assigning the uniquely mapped reads to genomic features through featureCounts (Liao et al. 2014).

DESeq2 was utilized to perform the Differential expression analysisby RStudio (version 4.4.2) (Love et al. 2014). The raw counts were first normalized through DESeq2 size factor normalization and transformed using the variance-stabilized transform (VST). Sample quality and consistency among biological replicates were assessed by hierarchical clustering based on a sample-to-sample distance matrix calculated from VST-transformed data. Samples identified as outliers or displaying inconsistent clustering relative to their biological replicates were excluded from subsequent analyses (Table S1). Principal component analysis (PCA) was performed on VST data. To maximize the resolution of biological variables beyond tissue-specific differences, the dataset was split into two different tissue-specific subsets. This approach allowed for the evaluation of Time while accounting for differences between natural and transplanted meadow plants without the primary effect driven by tissues effect. A likelihood ratio test (LRT) was employed on each dataset using a multi-factor design. For this test, the full model (∼ Time + Plant type) was tested against a reduced model excluding the Time term (∼ Plant type). The differentially expressed genes (DEGs) identified through this approach were filtered using false discovery rate (FDR) threshold of 0.001 and then clustered by using a hierarchical clustering. The identified cluster were represented through a Heatmap generated with the Pheatmap package (Kolde 2015)and the resulting dendrogram cut by using the Cutree function. A representation of the expression trend across the three time points of each cluster per dataset were made using lineplots made with the ggplot2 R package (Wickham 2011)

### 2.5 Co-expression network and functional analyses

VST data were used for the identification of co-expression modules by using the package Weighted Gene Co-expression Network Analysis (WGCNA). Modules were identified using dynamic tree cutting with a minimum size of 100 genes. For each module, module-trait associations were calculated as the Pearson correlation coefficient between the module eigengene (the principal component of the module expression profile) and the experimental conditions. Hub genes were defined as the 5% most connected genes within each module and co-expression network of these genes were made using Cytoscape where the connections were represented with edge and the degree (number of connections) through a gradient of color. Gene Ontology (GO) enrichment analysis was performed using the R package ClusterProfiler (Yu et al. 2012) with an adjusted p-value cutoff of 0.05 which was calculated using the Benjamini-Hochberg method. Results were visualized using Dotplots made in Rstudio with the package ggplot2.

## 3. Results

### 3.1 Response of *P. oceanica* seedlings to transplantation

At the time of transplantation, one-year old seedlings appeared healthy, with an average of 5 lush green leaves, up to 17 cm long (Figure 1A,B,G, Figure S1). Transfer from controlled ex situ conditions to the field was followed by a marked reduction in leaf number and total leaf length during the first month. Already one month after transplantation, the mean number of leaves dropped to 3.8, which reflected also on the total leaf length (Figure 1A,C). At later times, the number of leaves further decreased, and they shortened, likely as result of necrosis and herbivory (Figure 1A-C, Figure S1). Older leaves appeared brown, damaged and covered with epiphytes (Figure 1G). Conversely, the number of roots, but not their mean length, increased over time (Fig 1D-F). Natural meadows presented healthy bundles at all sampling times (Fig 1H).

**Figure 1.**
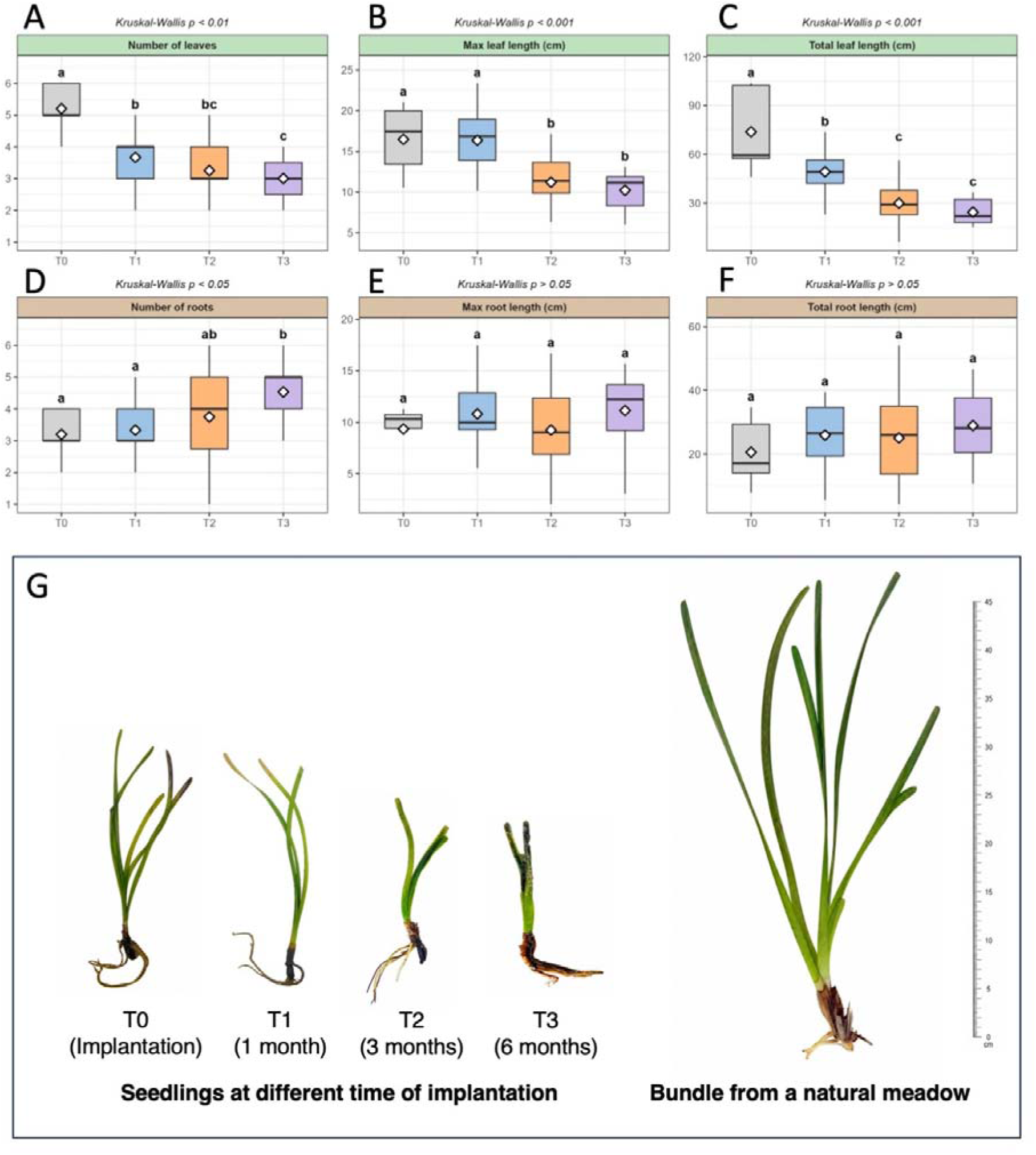
Leaf and root growth in transplanted seedlings. **A**: number of leaves. **B**: maximum leaf length; **C**: total leaf length; **D:** number of roots; **E**: maximum root length; **F**: total root length. All parameters were measured at transplantation (T0) and after one (T1), three (T2) and six months (T3). Different letters above the boxplots indicate a significant difference between time points (pairwise Mann-Whitney test, p < 0.05); time points sharing the same letter are not significantly different. **G**: representative seedlings at the different sampling times; **H**: representative bundle from a natural meadow at T2.

Analyses of water quality and microbiological load revealed consistent values in both transplantation sites, along time (Table S2), in agreement with the typical oligotrophic status of Mediterranean. Only on the Autumn sampling (T3), the bacterial load was higher and the total P content lower, at both sites, compared to previous measurements.

### 3.2 Temporal variation in gene expression in transplanted seedlings and natural meadow plants

A total of 1.1 billion paired-end (PE) reads were mapped to the reference genome, averaging 42 million reads per sample. Based on the sample-to-sample distance matrix analysis, eight (out of 72) seedling samples and three (out of 30) samples from natural meadows, mostly roots, were identified as outliers and excluded from subsequent analyses (Figure S4, Table S1).

The PCA performed on the complete transcriptional dataset revealed Tissue as the primary driver of the genetic variance, clearly segregating leaf and root samples into two different clusters along the first principal component (PC1), which accounted for 85% of the total variance (Figure 2A). To eliminate the confounding variance driven by tissue-specific gene expression and maximize the resolution of our experimental variables, the dataset was split into two independent subsets for roots and leaves. The PCA of leaf samples revealed a distinct distribution: PC1 (30% of the variance) explained the difference between transplanted seedlings and adult natural meadow plants, while PC2 (20%) captured the response to seasonality (Figure S5A). In roots, the relative contribution of these factors was even more skewed (51% for plant type and 14% for season, Figure S5B).

**Figure 2.**
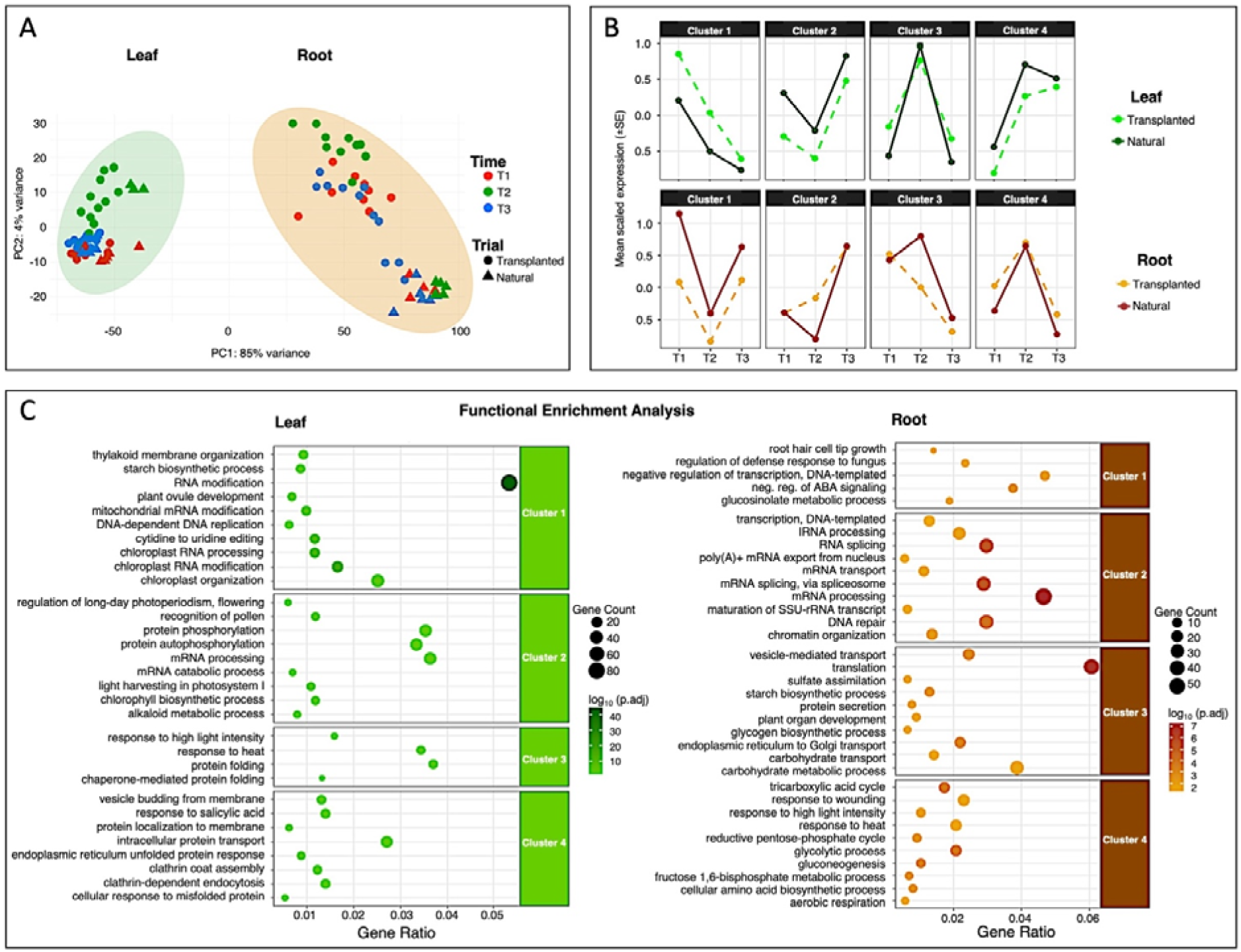
PCA and temporal expression analysis of *P. oceanica* tissues. **A** PCA plot of leaf and root samples showing variance distribution by tissue type. Colors and shapes indicate time points (T1– T3) and plant type (TRA, NAT). **B** Mean scaled expression trends across four clusters in leaf (green) and root (orange/red) tissues. Lines represent the temporal dynamics of gene expression for TRA and NAT plants. **C** Biological Process GO enrichment analysis of the identified clusters. Dotplots show the top enriched terms for leaf (green) and root (brown) clusters; dot size indicates gene count, whereas color represents significance.

Additional PCA analyses conducted to assess the similarity of the two transplantation sites on global gene expression patterns confirmed that samples randomly distributed between sites and groups within each site (Figure S6), in agreement with the chemical and microbiological equivalence of the two sites. Therefore, all biological replicates were treated together for subsequent analyses, regardless of site and group.

In order to identify DEGs with specific temporal trend, after the transplantation, we performed a LRT. The analysis led to the identification of 4564 DEGs for root and 6207 DEGs for leaf. A hierarchical clustering of these DEGs allowed the identification of genes cluster with similar trends across different time points (T1: spring; T2: summer; T3: autumn) (Figure S7). Four different clusters were identified for both root and leaf, ranging from 349 DEGs (cluster 4) to 1791 DEGs (cluster 1) for root and from 596 (cluster 3) to 2460 (cluster 1) for leaf (Tables S3, S4).

In root, cluster 1 showed a trend of up-regulation at both T1 and T3, whereas the opposite trend was observed in cluster 4 (Figure 2B). Clusters 2 and 3 showed a difference between transplanted seedlings and natural adult meadow plants. Cluster 2 was predominantly up-regulated at T3, but showed a reduction at T2 only in natural meadow plants. In cluster 3, the complementary peak at T2 was only evident in adult plants, whereas in seedlings there was a progressive reduction of expression along time. In leaves, the pattern of gene expression was similar between transplanted and natural meadow plants in all clusters. Cluster 1 showed a decreased expression along time, whereas cluster 4 had an opposite trend. Cluster 2 and Cluster 3 displayed mirror expression profiles, with Cluster 2 showing a sharp downregulation at T2, contrasting with the peak observed in Cluster 3.

Functional enrichment analysis of these clusters revealed the biological process in which DEGs were involved across the different seasons (Figure 2C). At T1, leaf DEGs were involved in RNA processing and modification and biosynthetic process of starch, while root DEGs were enriched in regulation of hormonal response (ABA) and transcription regulation. At T2, both root and leaf showed up-regulation of genes involved in response to heat and to high light intensity. Root DEGs were also enriched in regulation of carbohydrate and energy (glycolysis, tricarboxylic acid) metabolism, while leaf DEGs were involved in the regulation of protein folding process. At T3, root DEGs were mainly involved in the regulation of RNA processing, splicing and DNA repair, while leaf up-regulated DEGs were enriched in functions related to regulation of misfolded protein, response to salicylic acid and chlorophyll biosynthetic process.

### 3.3 Weighted gene co-expression network analysis and hub genes screening

To identify the co-expression modules correlated to the response of transplanted and natural meadow plants and the hub genes involved in their transcriptional regulatory network, a WGCNA was carried out, including the whole dataset of 23,306 genes for leaf and root. The analysis revealed 21 co-expressed modules in the leaf and 17 in the root, ranging from 174 to 1848 and from 341 to 3,277 genes, respectively (Figure S8). To relate co-expression modules to specific sample conditions, Pearson correlation coefficients were calculated using module eigengenes (MEs) (Figure 3A).

**Figure 3.**
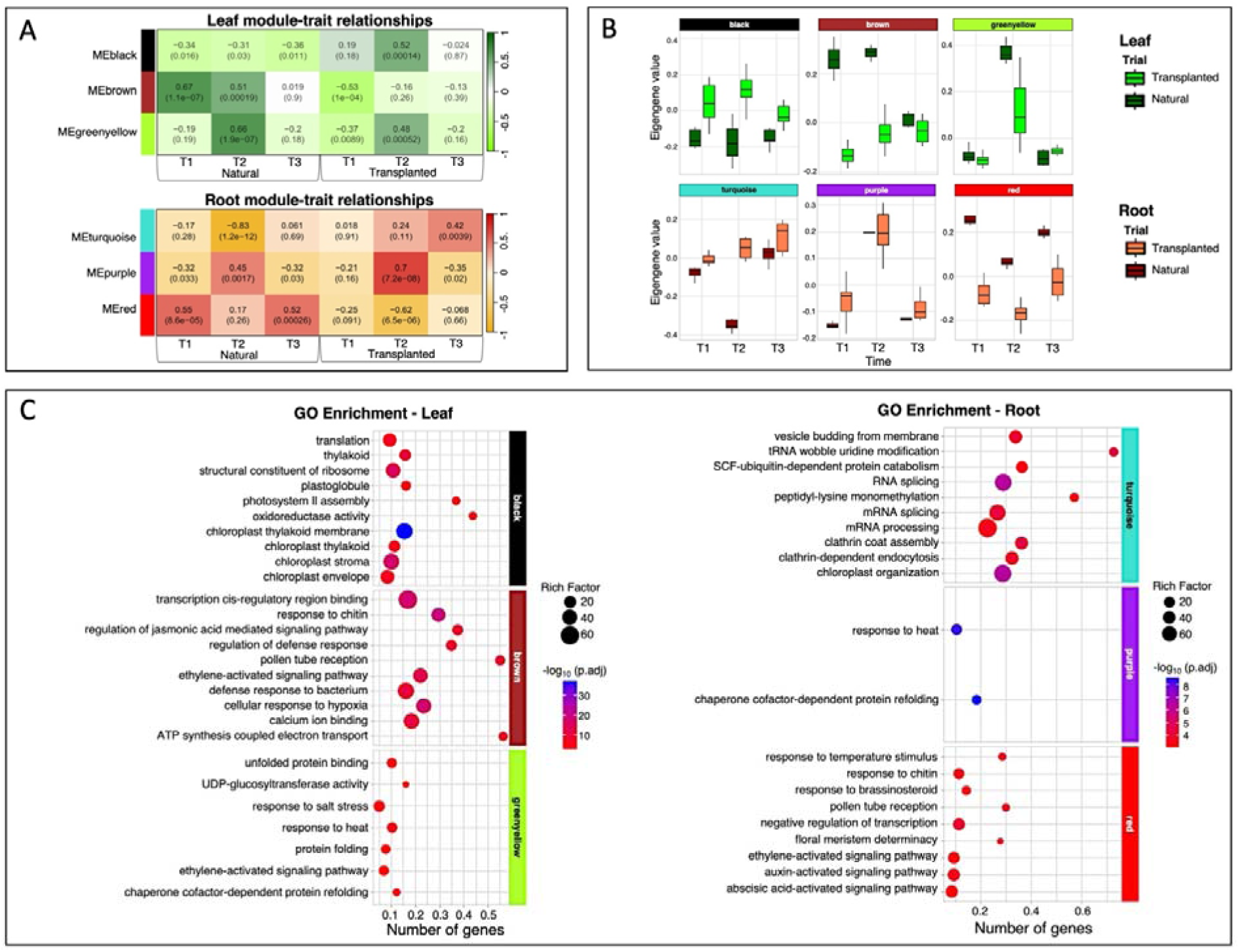
WGNA of *P. oceanica* samples. **A** Module-trait associations revealed by the Pearson correlation coefficient for leaf and root tissues. The leftmost color column indicates co-expression modules. Numbers indicate correlation coefficients within each time point andtplant type. **B** Module eigengene temporal trends (T1–T3) across the selected WGCNA modules for transplanted (TRA) and natural (NAT) plants. **C** Biological Process GO enrichment analysis of the corresponding modules. Dotplots show enriched terms for leaf-specific (“black”, “brown”, “greenyellow”) and root-specific (“turquoise”, “purple”, “red”) modules; dot size indicates the number of genes whereas color represents the significance.

Significant differences were observed for the leaf “brown” and root “red” modules, which maintained higher expression levels in natural *P. oceanica* meadow plants than in transplanted seedlings over time (Figure 3B). Specifically, the leaf “brown” module comprised genes involved in defense responses, ATP synthesis, and the regulation of hormone pathways such as jasmonic acid and ethylene, while the root “red” module was tied to the regulation of responses to ethylene, auxin and ABA (Figure 3C). In contrast, the leaf “greenyellow” and root “purple” modules showed a marked induction specifically at T2, displaying higher eigengene values in seedlings than in adult plants. These two co-expressed modules were both enriched in functions related to the regulation of protein folding and heat response. A different temporal trend was observed for the root “turquoise” module, which reached its maximum expression levels at T3 in both plant types despite a significant negative correlation recorded for natural meadow plants at T2. This module was characterized by functions related to RNA processing and DNA repair mechanisms. Finally, the leaf “black” module displayed a distinct profile, showing a specific positive correlation with the transplanted seedlings at all time points while remaining largely down-regulated or stable in natural meadow plants; genes within this module were exclusively involved in the regulation of photosynthesis.

The identification of hub genes provided further resolution into the transcriptional network of these modules, as visualized in the network plots showing the highest connectivity for nodes (Figure 4, Table S5). In the leaf, the “brown” module maintained higher expression levels in natural meadow plants than in transplanted seedlings over time with 50 hub genes detected. The highest connectivity was observed for several transcription factors (TFs), specifically from the WRKY and ERF (Ethylene Response Factor) families, along with genes involved in the jasmonic acid biosynthetic pathway, such as *lipoxygenase 2* and *allene oxide cyclase 3*, and several genes involved in the synthesis and modification of the cell wall, such as *cellulose synthase* and *pectine methylesterase*. Similarly, the root “red” module, which also displayed consistently higher expression levels in natural meadow plants with 23 hub genes, was characterized by central regulators linking cytokinin biosynthesis and long-distance transport with root developmental programs and environmental signal perception. Hub genes included those encoding for the cytokinin biosynthetic enzyme AtIPT3 and the cytokinin exporter ABCG14, together with several transcriptional regulators such as homeobox-leucine zipper protein HAT14, MYB61, and TCP domain protein 4. For the modules induced at T2, the “greenyellow” (leaf; 17 hub genes) and “purple” (root; 23 hub genes) hubs were predominantly composed of heat shock proteins (HSPs), such as HSP70, HSP90, and HSP 22 kDa protein, various chaperones (e.g., DnaJ) and a TF involved in heat stress response (*Heat stress transcription factor 24*). In the root “turquoise” module, which reached its maximum expression levels at T3 in both plant types, the 109 hub genes were primarily represented by genes involved in the regulation of signaling process like *serine/threonine-protein kinase 10* or *dual specificity protein phosphatase 12,* by components of the RNA splicing and processing such as *pre-mrnaprocessing factor 19* and *rna polymerase sigma factor sigma* and DNA repair enzymes such as *dna-binding transcription factor 2*. Finally, the leaf “black” module, which showed a specific positive correlation with the transplanted seedlings across all time points, included hub genes encoding structural components of the photosystem II (PSII) like *Protein maintenance of PSII under high light*, *Photosystem II 10 kDa polypeptide* and *Light-Harvesting Complex II,* and protein involved in photo and stress protection like *violaxanthin de-epoxidase* and *stress enhanced protein 1*.

**Figure 4.**
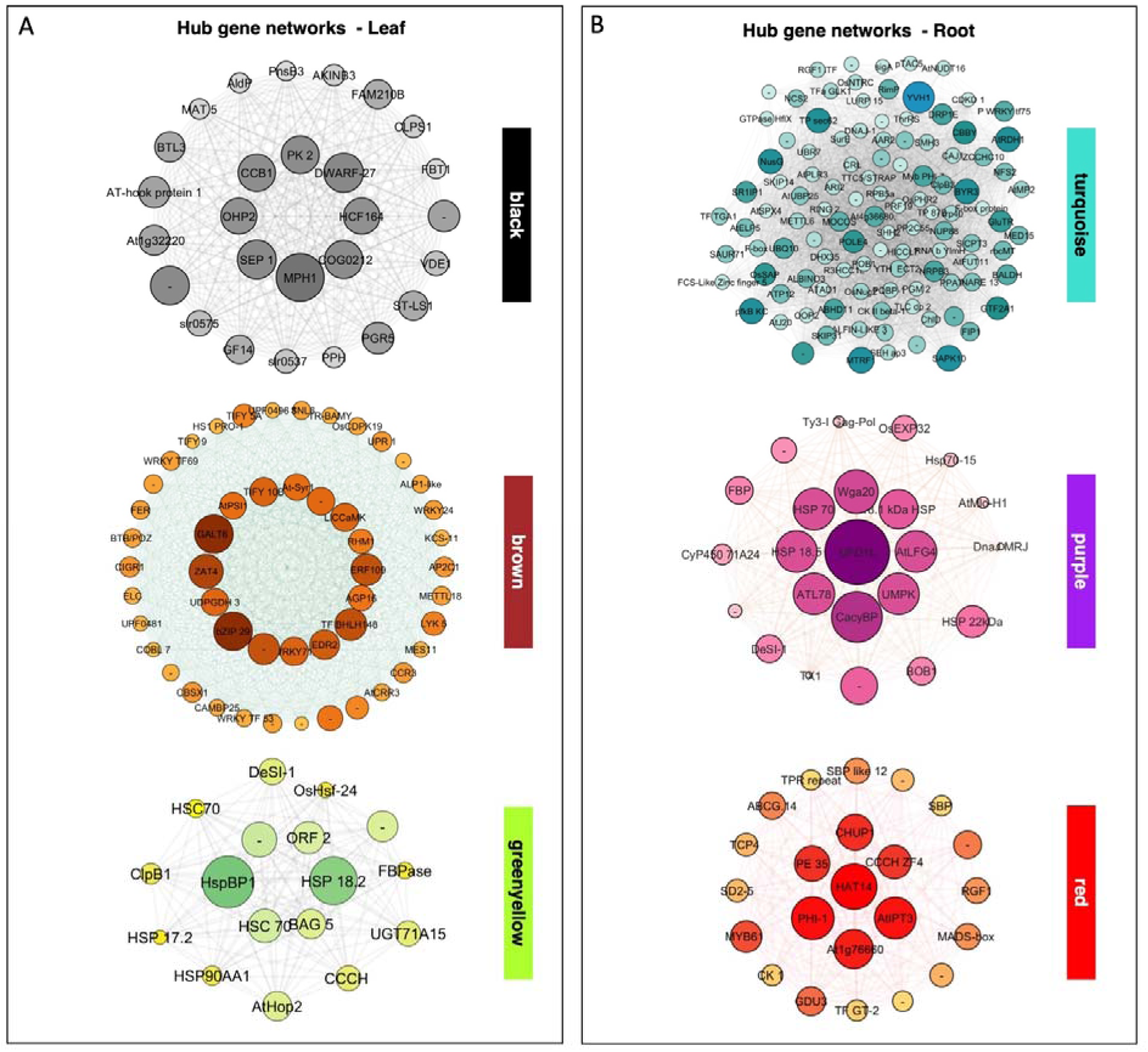
Hub gene network analysis within the identified co-expression modules. **A** Gene interaction networks for the modules associated with leaf tissues (“black”, “brown”, “greenyellow”). Nodes represent individual genes, and edges indicate co-expression strength. Central, larger nodes identify the top hub genes within each network. **B** Hub gene networks for the modules associated with root tissues (“turquoise”, “purple”, “red”). The intensity of node color and size reflect the degree of connectivity (intramodular connectivity) and significance within the module, highlighting key regulatory candidates for tissue-specific responses.

## 4. Discussion

### 4.1 Transcriptional differences between transplanted seedlings and natural meadow plants

The primary objective of this study was to characterize the transcriptional landscape of *P. oceanica* seedlings during the critical establishment phase following transplantation into a natural environment. By comparing the molecular signatures of transplanted seedlings and established adult meadow plants across tissues and sampling periods, we aimed to identify the transcriptional programs underlying early adaptation and to evaluate the molecular consequences of restoration practices.

The global PCA revealed a clear separation between leaf and root samples, confirming that tissue identity is the primary determinant of transcriptomic variation in *P. oceanica* (Valenti et al. 2026), irrespective of plant type or sampling time. Such tissue-specific differentiation reflects the functional specialization of seagrasses, with leaves primarily devoted to photosynthetic carbon assimilation, as previously described for *Zostera marina* (Olsen et al. 2012), whereas roots are specialized for anchorage and belowground resource acquisition, consistent with developmental transcriptomic profiles reported for *P. oceanica* (Valenti et al. 2026).

When tissues were analysed separately, PCA provided a higher-resolution view of plant type-specific transcriptomic variation across seasons. In both leaves and roots, natural and transplanted samples remained significantly separated along PC1 throughout the experimental period, indicating that transplanted seedlings retained a distinct transcriptional identity even after several months of exposure to natural environmental conditions. A likely explanation for this persistent divergence is the difference in developmental stage between the two plant types. Transplanted individuals were young seedlings (12–18 months old), whereas natural samples originated from mature meadow plants. Developmental stage has recently been shown to exert a strong influence on gene expression in *P. oceanica* seedlings (Valenti et al. 2026), and young plants activate specific transcriptional and protective pathways during environmental acclimatisation, including responses to thermal stress (Pazzaglia et al. 2025). A second, non-mutually exclusive explanation is that transplantation itself induces a prolonged transcriptional reprogramming. Seedlings were transferred from stable, controlled laboratory conditions to a highly dynamic natural environment characterized by fluctuations in temperature, light, hydrodynamics and sediment chemistry. Such environmental transitions are expected to require extensive physiological adjustment during acclimatisation to field conditions. Additionally, the very act of transplantation has long lasting consequences, since leaf length of cutting does not recover within six years after transplant (Pansini et al. 2024). Consequently, the transcriptomic divergence observed here may reflect the combined effects of ontogeny, previous growth under controlled conditions and acclimatisation to the field environment.

The persistent separation between the groups should therefore not be interpreted solely as evidence that transplanted seedlings had failed to reach, or were recovering towards, the molecular state of established adult plants. Although the approximately one-year age of the seedlings resulted from the prolonged ex situ maintenance preceding the spring transplantation, it also allowed field acclimatisation to be examined beyond the earliest developmental phase, when growth is strongly supported by the mobilisation of seed-derived nutrient reserves. Notably, transcriptomic divergence between natural and transplanted plants was more pronounced in roots than in leaves, suggesting that belowground tissues are particularly sensitive during the post-transplantation phase. Root systems are directly exposed to local sediment properties, including oxygen availability, nutrient gradients and microbial communities, all of which are likely to influence gene expression. Differences in substrate composition between transplanted gravel and the natural matte, together with the progressive establishment of root-associated bacterial communities (Boulenger et al. 2025), may therefore contribute to the persistent transcriptional differences observed between plant types. Collectively, these findings indicate that the transcriptional profile of transplanted seedlings does not immediately reflect the molecular state of natural meadows but instead initiates a gradual process of transcriptional reprogramming, with belowground tissues apparently requiring a longer period to reach functional convergence.

### 4.2 Seasonal transcriptomic plasticity of *P. oceanica*

The second principal component of the PCA captured the seasonal component of transcriptomic variation and revealed that mature meadows and transplanted plants maintained distinct transcriptional trajectories throughout the study period. The persistent separation between the two plant types indicates that transplanted seedlings do not simply mirror the seasonal responses of established meadows but instead exhibit type-specific transcriptional adjustments to the changing environmental conditions.

The LRT identified the subset of genes whose expression was significantly modulated over time, highlighting temporal changes in both leaf and root transcriptomes during the six-month post-transplantation period. At T1 (late May, one month after transplantation), transplanted seedlings exhibited a pronounced induction of genes involved in RNA processing and RNA modification, particularly in leaves. Similar transcriptional responses have been reported in the seagrass *Cymodocea nodosa* under multiple abiotic stress conditions (Malandrakis et al. 2017), suggesting that extensive post-transcriptional regulation represents an early component of stress acclimatisation in marine angiosperms. Consistent with previous observations in *P. oceanica* (Dattolo et al. 2014), this transcriptional reorganization likely reflects the rapid adjustment of regulatory networks following the transition from controlled laboratory conditions to a highly variable natural environment.

In parallel, roots showed a selective enrichment of genes associated with ABA signalling. ABA is a central regulator of osmotic homeostasis and stress-induced transcriptional reprogramming in terrestrial angiosperms (Cutler et al. 2010), and its activation in transplanted seedlings suggests that belowground tissues rapidly perceive and respond to changes in sediment conditions following transplantation. Together, these coordinated responses indicate that the first month after transplantation is characterized by extensive molecular reorganization aimed at establishing regulatory homeostasis in both above- and belowground tissues.

By T2 (late July, three months after transplantation), we observed a coordinated induction of genes involved in heat response and protein folding, consistent with the prolonged high seawater temperatures recorded during summer. Similar activation of heat-responsive pathways has been described in *Z. marina* exposed to elevated temperatures (Franssen et al. 2011), suggesting that these mechanisms are broadly conserved among seagrasses. Importantly, the induction of these pathways occurred in both mature meadows and transplanted plants, indicating that the shared response in both groups is consistent with exposure to the same summer thermal conditions.

This transcriptional response coincided with the regional marine heatwave documented in the western Mediterranean during summer 2022, when estimated seawater temperature at the study side reached approximately 28 °C (Figure S3). In Western Mediterranean, the summer 2022 anomaly peaked the record figure of 4.6 °C above the average of the past four decades (Marullo et al. 2023; Astruch et al. 2026). The severity of the event was further supported by field observations of visible stress in gorgonian populations inhabiting deeper areas of the study site (as reported by the diving service) and other parts of the Mediterranean (Estaque et al. 2023). Such extreme thermal events are becoming increasingly frequent in the Mediterranean Sea, where seawater temperature has increased by approximately 0.041 °C year ¹ (Copernicus Marine Service), contributing to a higher incidence of marine heatwaves (Garrabou et al. 2022). Understanding how both natural meadows and restored *P. oceanica* plants respond to these increasingly common events is therefore essential for predicting the long-term resilience of Mediterranean seagrass ecosystems.

Previous studies have associated widespread mortality of *P. oceanica* meadows with the 2003 and 2006 Mediterranean heatwaves, during which seawater temperatures also approached 28 °C (Díaz-Almela et al. 2009; Marbà and Duarte 2010). Experimental studies have shown that *P. oceanica* seedlings achieve maximum gross photosynthetic performance at approximately 28 °C under short-term exposure (Rinaldi et al. 2023). However, these experiments evaluated acute thermal stress over 24 hours, whereas natural marine heatwaves often persist for several weeks or months. Indeed, prolonged exposure to temperatures above 25 °C has previously been shown to reduce seedling growth (Olsen et al. 2012). In the present study, seawater temperatures remained close to or above 28 °C for nearly two months (Figure S3), exposing plants to chronic rather than acute thermal stress. Consistent with these environmental conditions, seedlings collected at T2 and T3 displayed reduced leaf growth and extensive necrotic areas, indicating that prolonged exposure imposed a substantial physiological burden.

The molecular responses observed at T2 were consistent with this prolonged thermal stress. In roots, differentially expressed genes were enriched for glycolysis, the tricarboxylic acid cycle and other pathways involved in carbohydrate and energy metabolism, suggesting an increased allocation of metabolic resources to sustain the energetic demands imposed by elevated temperature. Similar metabolic adjustments have been reported in heat-stressed seagrasses (Marín-Guirao et al. 2017). In leaves, the enrichment of protein folding and molecular chaperone pathways likely reflects the activation of proteostasis mechanisms required to prevent protein denaturation and maintain cellular function under prolonged heat stress, consistent with previous observations in *Z. marina* (Franssen et al. 2011).

By the final sampling point (T3, late October), transcriptomic regulation shifted from stress responses towards processes associated with cellular maintenance and recovery. In roots, differentially expressed genes were enriched for RNA processing, mRNA splicing and DNA repair, whereas leaves showed enrichment of genes involved in protein quality control and salicylic acid signalling, processes widely associated with stress recovery and cellular protection (Filgueiras et al. 2019). Rather than indicating that plants were still experiencing severe stress, the activation of DNA repair pathways may reflect recovery from the cumulative molecular damage incurred during the summer heatwave. This interpretation is consistent with the “heat stress memory” described by (Pazzaglia et al. 2025), in which *P. oceanica* seedlings activate genome maintenance and repair mechanisms following thermal stress to restore transcriptional homeostasis. The seasonal timing of this response is also noteworthy. Autumn marks the transition towards reduced metabolic activity in temperate seagrasses, and the transcriptomic stabilization observed here may therefore represent the combined effects of recovery from summer stress and the onset of seasonal metabolic downregulation.

### 4.3 Co-expression modules differ between tissues and plant type

The WGCNA analysis provided complementary insights into the transcriptional differences between natural meadow plants and transplanted plants by identifying coordinated gene networks rather than individual differentially expressed genes. Overall, the results indicate that restoration status influences not only the expression of specific genes but also the organization and stability of transcriptional regulatory networks. Several co-expression modules displayed consistent type-specific expression patterns throughout the experiment, suggesting that transcriptional differences between transplanted seedlings and established adult plants persisted over the six-month study period.

In leaves, the higher expression of the “brown” module in natural meadows suggests that long-established meadows maintain a more stable transcriptional program associated with structural integrity and defence. This module includes genes involved in cell wall organization, such as *pectin methylesterase*, together with components of jasmonic acid signalling, including *lipoxygenase 2* and *allene oxide cyclase 3*. These pathways contribute to cell wall remodelling, defence signalling and tissue integrity (Wolf 2017; Wasternack and Song 2017), processes that are likely to support the long-term stability of mature leaves. The central position of WRKY transcription factors and cellulose synthase within this module further supports this interpretation. WRKY proteins are widely recognized as master regulators of plant stress-responsive transcriptional networks (Wang et al. 2011), whereas cellulose synthase is essential for maintaining cell wall architecture and mechanical stability (Wolf 2017). Interestingly, expression of the leaf “brown” module progressively converged between transplanted and natural meadow plants by T3, suggesting that transplanted seedlings gradually acquire components of the regulatory network characteristic of established meadows.

In contrast, plant type-specific differences remained more pronounced in roots. The “red” module exhibited consistently higher expression in natural meadows plants throughout the study, indicating that root module expression remained more distinct between the two plant types over the experimental period. Among the central hub genes of this module are *ABCG14*, which mediates the root-to-shoot transport of cytokinins (Ko et al. 2014), and *HAT14*, a homeodomain transcription factor involved in root development and environmental adaptation (Chew et al. 2013). These hubs suggest that mature root systems possess well-established hormonal and developmental regulatory circuits that are not yet fully established in recently transplanted seedlings. Consequently, although leaf transcriptomes showed evidence of progressive convergence, root tissues appeared to retain a less mature regulatory organization throughout the experimental period.

These contrasting tissue-specific dynamics emphasize that temporal changes following transplantation were not uniform across the plant. Leaf module expression showed greater similarity between groups by the final sampling point, whereas differences in root-associated modules persisted throughout the study. Because root contribute to nutrient acquisition, anchorage and interactions with sediment microbial communities, the delayed establishment of these regulatory networks may represent one of the principal bottlenecks limiting the functional integration of restored seedlings. More broadly, molecular profiles may complement traditional descriptors such as survival and growth in future evaluations of transplantation outcomes.

### 4.4 Adaptive transcriptional strategies underlying seasonal acclimatisation in *P. oceanica*

The temporal dynamics of the WGCNA modules demonstrate that seasonal environmental variation, particularly the summer temperature peak, reshaped gene co-expression networks in both natural meadow plants and transplanted seedlings. Rather than representing isolated transcriptional responses, these modules revealed coordinated regulatory programs that underpin the seasonal acclimatisation of *P. oceanica*. Although both plant types activated common stress-response pathways, differences in the magnitude and persistence of module expression indicate that transplanted seedlings and established meadow plants adopt partially distinct strategies to cope with environmental fluctuations.

The most evident example of this coordinated response is represented by the leaf “greenyellow” and root “purple” modules, both of which reached maximal expression during T2. These modules were enriched in heat-responsive genes, including *HSP70*, *HSP90* and several additional molecular chaperones, highlighting the central role of proteostasis in maintaining cellular function during prolonged thermal stress. Heat shock proteins are among the most conserved components of the plant stress response and prevent protein denaturation, aggregation and irreversible damage under elevated temperatures. Their induction during the summer heatwave is therefore consistent with previous studies demonstrating that marine heatwaves trigger extensive chaperone-mediated responses in *P. oceanica*, with the magnitude of activation varying among populations originating from different thermal environments (Stipcich et al. 2024).

Interestingly, the expression of the leaf “greenyellow” module was consistently higher in natural meadow plants, whereas the root “purple” module showed comparable expression between plant types. These patterns suggest that the core molecular machinery responsible for thermal protection is largely conserved across the meadow plant, supporting previous physiological observations that both natural meadow plants and restored plants activate similar protective mechanisms during periods of elevated temperature (Marín-Guirao et al. 2017, 2018). At the same time, the stronger induction observed in leaves of natural meadow plants may indicate a more efficient or more tightly coordinated transcriptional response in established individuals, reflecting the progressive optimization of regulatory networks during meadow maturation rather than the activation of fundamentally different pathways.

In contrast, the leaf “black” module displayed a markedly different temporal behaviour, remaining consistently more highly expressed in transplanted plants throughout the experiment. This module comprised genes involved in photosystem II maintenance, photoprotection and repair, including *violaxanthin de-epoxidase*, PSII repair proteins, the 10-kDa PSII polypeptide and light-harvesting complex II components. The sustained activation of these pathways suggests that transplanted seedlings continuously invest resources in maintaining photosynthetic performance under fluctuating environmental conditions. Photosynthetic repair and photoprotection are tightly linked to the detoxification of reactive oxygen species generated under stress (Jahns and Holzwarth 2012), indicating that transplanted seedlings have a greater requirement for maintaining photosystem integrity than plants growing within established meadows. Rather than representing an acute stress response, the persistent activation of this module throughout the study suggests that recently transplanted seedlings sustain a chronically higher metabolic investment in photosynthetic maintenance while acclimating to the natural environment.

This molecular signature is consistent with field observations. Throughout the study, transplanted seedlings exhibited visible symptoms of physiological stress, including reduced leaf growth and necrotic tissues, likely resulting from the combined effects of transplantation, prolonged thermal stress, herbivory and the increased environmental variability experienced outside controlled laboratory conditions. In contrast, shoots collected from the natural meadow plant showed no evident signs of damage. Similar physiological resilience has previously been described for adult *P. oceanica*, particularly in populations naturally exposed to warmer environments, which can maintain photosynthetic homeostasis despite elevated seawater temperatures (Olsen et al. 2012; Marín-Guirao et al. 2017, 2018). Together, these observations suggest that natural meadow plants possess regulatory networks that minimize the energetic costs associated with prolonged stress, whereas recently transplanted seedlings must allocate a greater proportion of their resources to cellular maintenance and repair.

The progressive stabilization of the transcriptome became more evident by the final sampling point. In leaves, the convergence of eigengene expression within the “brown” module between natural meadow plants and transplanted plants suggests that transplanted seedlings gradually reconstructed regulatory programmes associated with tissue organization and defence. In roots, the “turquoise” module, enriched in genes involved in RNA splicing and genome maintenance, including the spliceosomal component *pre-mRNA-processing factor 19*, reached its highest expression at T3 in both plant types. *PRP19* plays a dual role in pre-mRNA processing and DNA repair, contributing to both transcriptome integrity and genome stability (Chanarat and Sträßer 2013). Interestingly, the temporary reduction of this module in natural meadow plants during T2 suggests that mature plants transiently down-regulated maintenance processes during periods of peak thermal stress, potentially reallocating metabolic resources towards immediate protective functions before restoring housekeeping activities once environmental conditions improved. In contrast, transplanted seedlings maintained a more sustained activation of these processes throughout the season, consistent with the continued need to support both environmental acclimatisation and developmental maturation.

Overall, the network-level analyses indicate that acclimatisation of *P. oceanica* seedlings was not achieved through the activation of novel stress-response pathways, but rather through the progressive reorganization and stabilization of existing regulatory networks. While natural meadow plants and transplanted plants shared the same core molecular mechanisms of seasonal acclimatisation, recently transplanted seedlings appeared to incur higher energetic costs associated with photosynthetic maintenance, cellular repair and regulatory reprogramming. These findings identify coordinated transcriptional processes potentially associated with the field acclimatisation of transplanted seedlings. Determining whether these processes contribute to successful establishment will require longer-term monitoring linking molecular patterns to independent ecological outcomes, including survival, rooting, growth and vegetative expansion.

### 4.5 Conclusions

This study provides the first comprehensive characterization of the seasonal transcriptomic dynamics of *Posidonia oceanica* seedlings during their establishment in a natural environment following transplantation. By integrating differential expression and gene co-expression network analyses across tissues and seasons, we show that transplanted seedlings undergo substantial transcriptomic reorganization during field acclimatisation while remaining distinct from established meadow plants, particularly in roots.

Both transplanted seedlings and natural meadow plants showed induction of heat-response and protein-folding pathways during the period of elevated summer temperatures, indicating that they shared core molecular responses to seasonal thermal conditions. However, transplanted seedlings consistently maintained higher expression of genes associated with photosystem II maintenance, photoprotection, protein turnover and cellular repair. Together with the observed reduction in leaf growth and the occurrence of necrotic tissue, these transcriptional patterns are consistent with a greater requirement for photosynthetic maintenance and protection during acclimatisation to field conditions, although the associated physiological and energetic costs were not directly measured.

Temporal responses differed markedly between tissues. The expression profiles of several leaf co-expression modules became progressively more similar between transplanted seedlings and natural meadow plants by the final sampling, suggesting partial convergence of leaf regulatory networks during acclimatisation. In contrast, differences in root-associated co-expression modules persisted throughout the study. Given the central roles of roots in anchorage, nutrient acquisition and interactions with sediment-associated microbial communities, these persistent transcriptomic differences may be relevant to longer-term seedling performance. However, establishing their ecological significance will require longer-term monitoring linking molecular patterns with independent measures of survival, rooting, growth and vegetative expansion.

More broadly, our findings demonstrate how transcriptomic analyses can complement traditional ecological descriptors by revealing molecular responses accompanying the transition from controlled cultivation to natural field conditions. The differentially expressed genes and co-expression modules identified here represent candidate molecular indicators of post-transplant acclimatisation and responses to prolonged summer warming, rather than direct indicators of restoration success or thermal resilience. As marine heatwaves become increasingly frequent under climate change, understanding the molecular processes associated with acclimatisation will improve our ability to interpret plant responses during restoration and to develop mechanistic frameworks for evaluating restoration trajectories. Future studies integrating transcriptomic, physiological, microbial and ecological measurements over longer temporal scales will be essential to determine whether these molecular patterns predict successful field establishment and long-term restoration outcomes for seagrass species.

## Authors’ contributions

FB, RDM: conceptualisation; GP, RDM: data curation; GV, FCo, GP, RDM: formal analysis; FB, FCa, RDM: funding acquisition; AS, VMG, RDM: investigation; VMG, GP, RDM: methodology; FCa, FB, RDM: project administration; AS, RDM: resources; GP, RDM: supervision; GV, RDM: validation; GV, FCo, FCa, RDM: visualisation; GV, RDM: writing-original draft; all authors: writing-review and editing.

## Acknowledgments

This work was supported by the Ministry of Education, University and Research (MUR), Italy [project Marine Hazard, grant number PON03PE_00203_1].

We are grateful to Arturo Zenone (Anton Dorhn Zoological Station, Palermo, Italy) and Marco Martinez (CNR IRBIM, Palermo, Italy) for assistance in seedling potting. We are also grateful to Gaspare Buffa and Carlo Patti (CNR IRBIM) and Debora Giampani (LUNA BLU diving) for assistance in seedling transplant.

## Supplementary data

**Figure S1.**
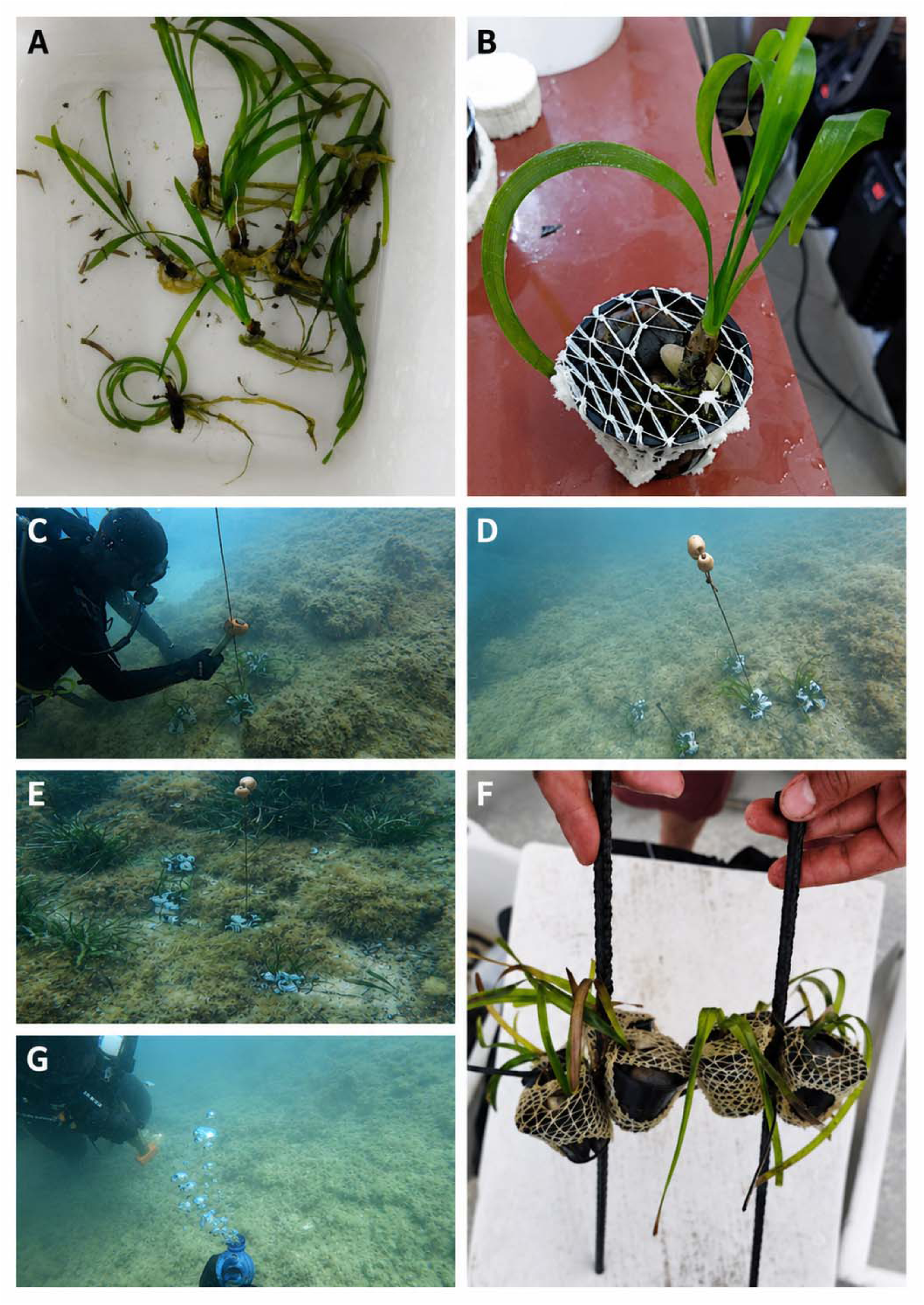
Transplantation of seedlings. **A**: seedlings ready to be potted; **B**: seedling in a pot with pebble stones, covered by elastic net; **C-E**: rebars transplanted in groups and signaled by buoys; **F**: rebars with seedlings at T1; **G**: water sampling.

**Figure S2.**
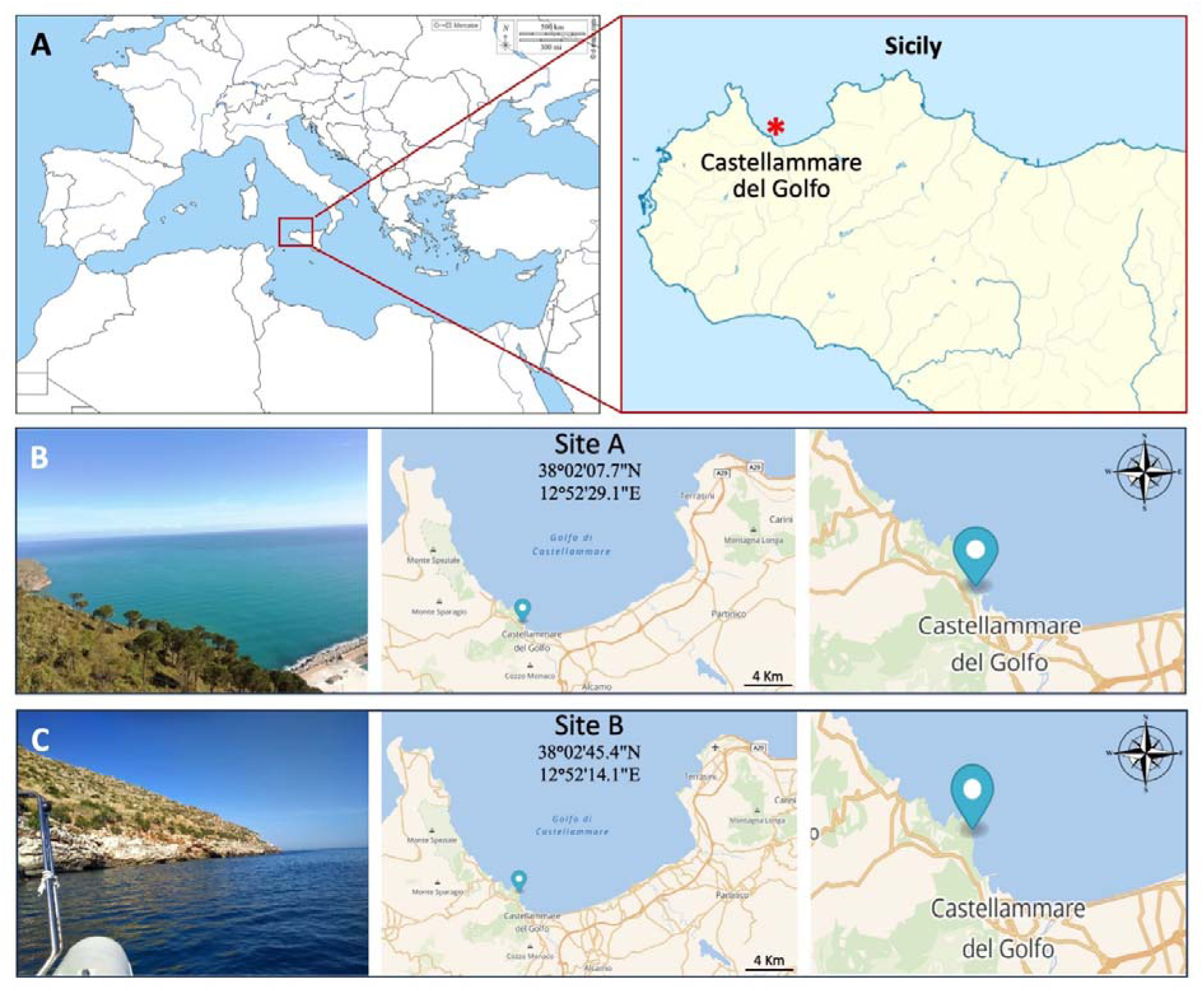
Transplantation site. **A**: map of the transplantation site; **B, C**: views of the site.

**Figure S3.**
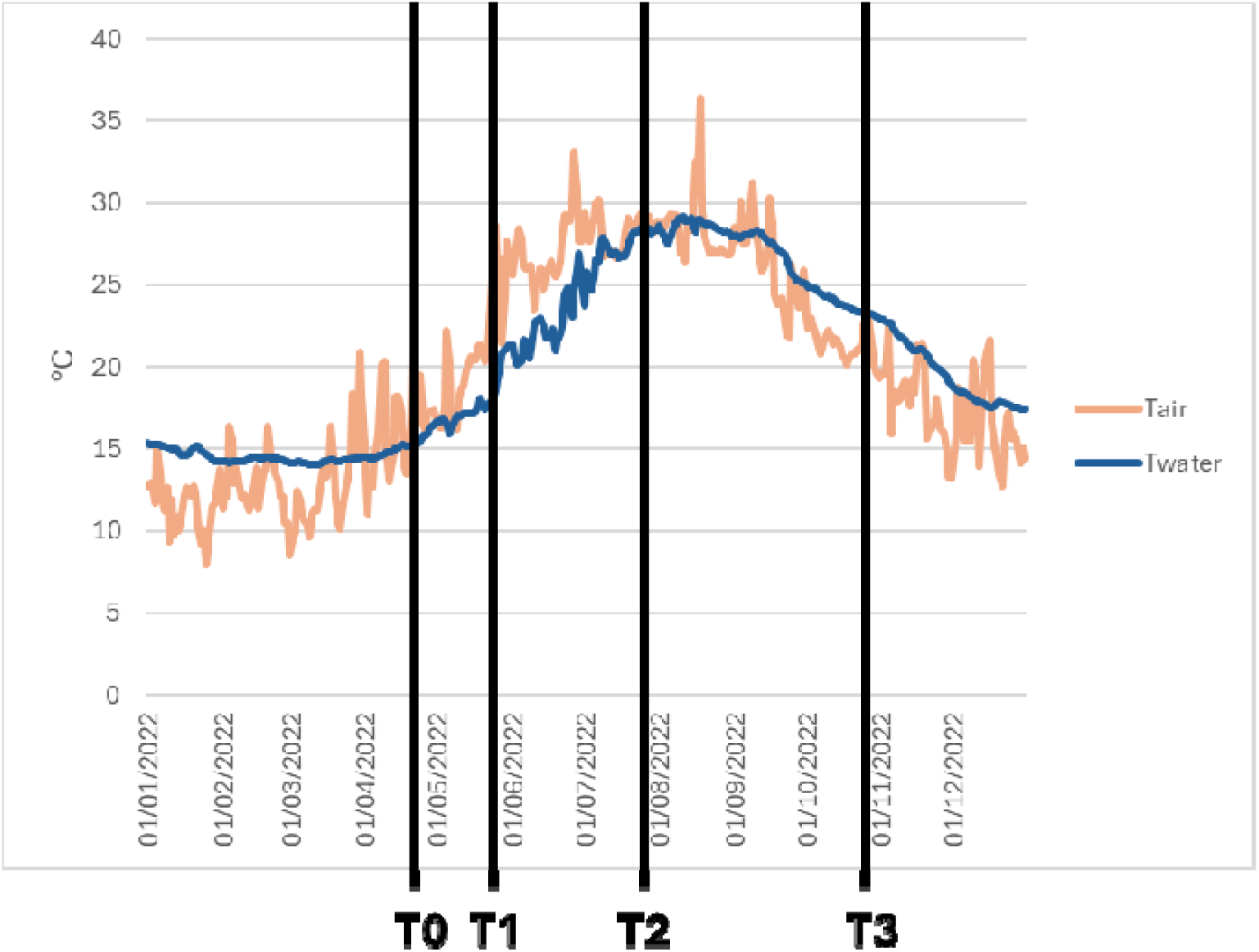
Daily mean air and seawater (5.46 m depth) temperatures in 2022. Transplantation (T0) and collection times (T1-T3) are indicated.

**Figure S4.**
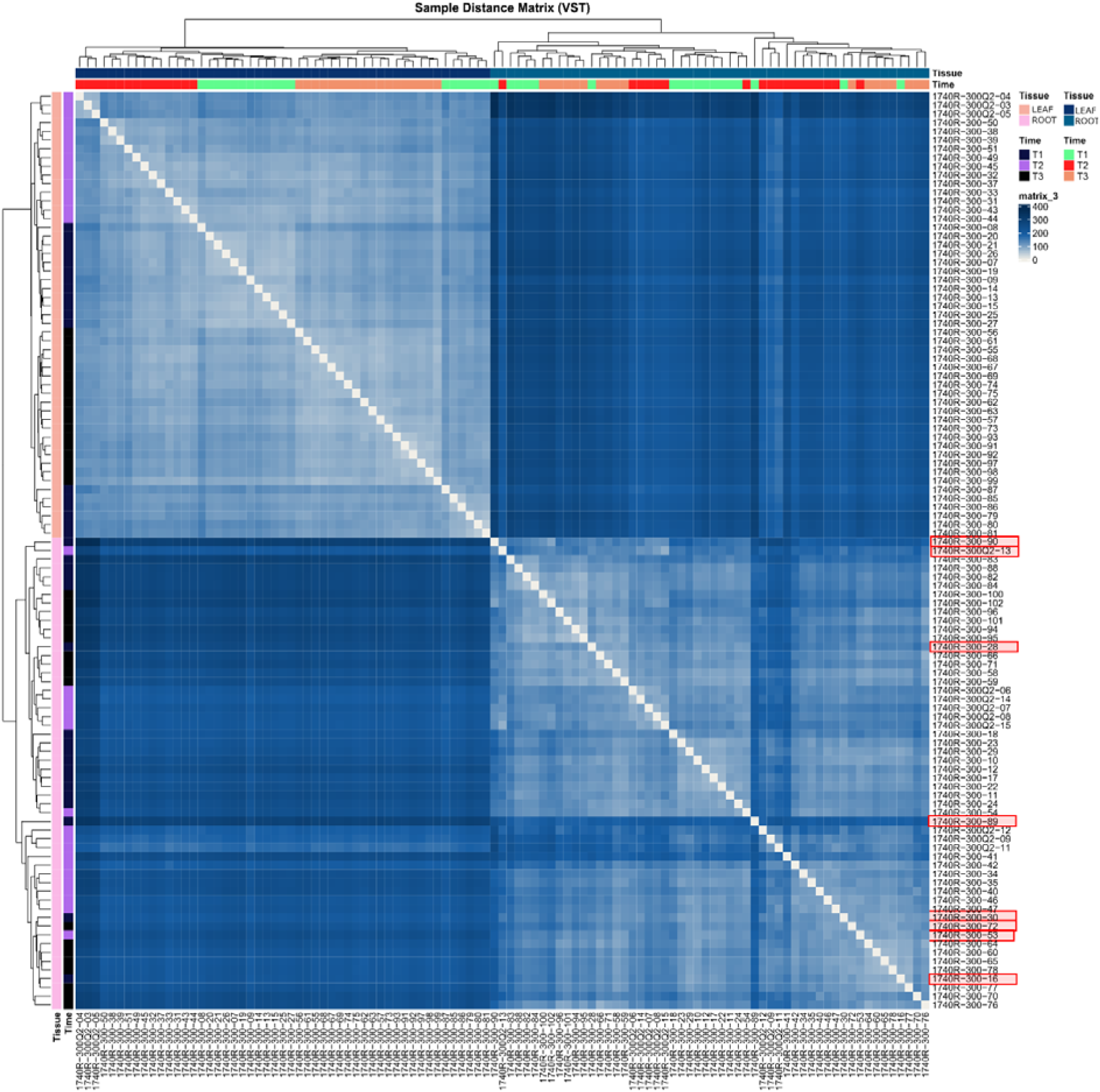
Sample-to-sample distance matrix based on VST gene expression data. Hierarchical clustering revealed a clear separation between leaf and root samples, indicating tissue-specific transcriptional profiles. Samples highlighted in red were identified as outliers because they did not cluster consistently with their corresponding biological replicates and were therefore excluded from downstream analyses.

**Figure S5.**
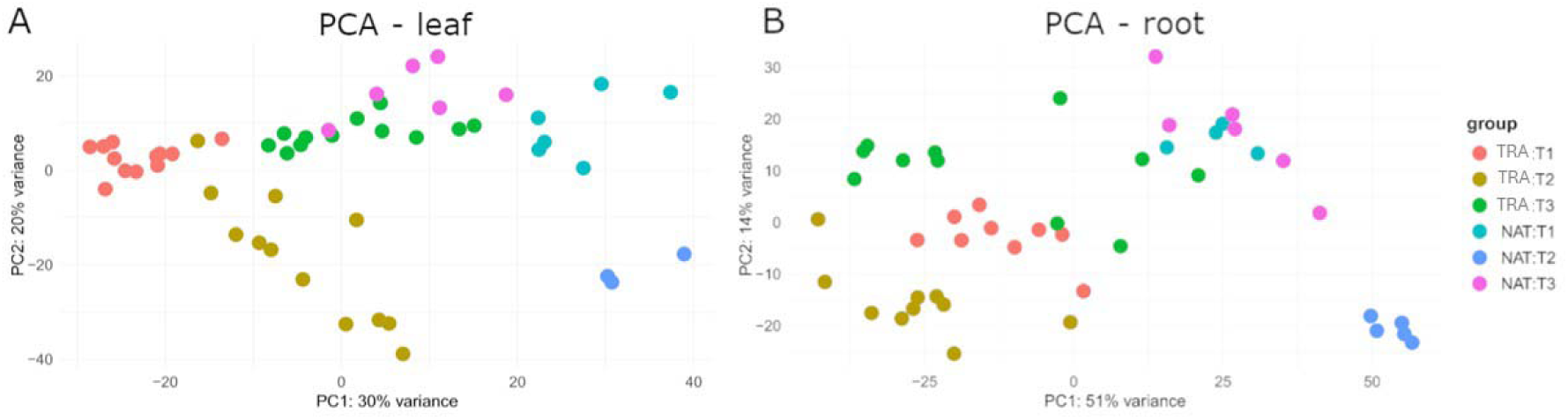
PCA of *P. oceanica* transcriptomes within specific tissues. **A** PCA plot of leaf samples which shows the separation of samples based on time points (T1–T3) and Plant types (TRA, NAT). **B** PCA plot of root samples which highlight the variance distribution, showing a distinct clustering of NAT and TRA plants across the three sampling time points. Colors represent the experimental groups as defined in the legend (Plant type:Time).

**Figure S6.**
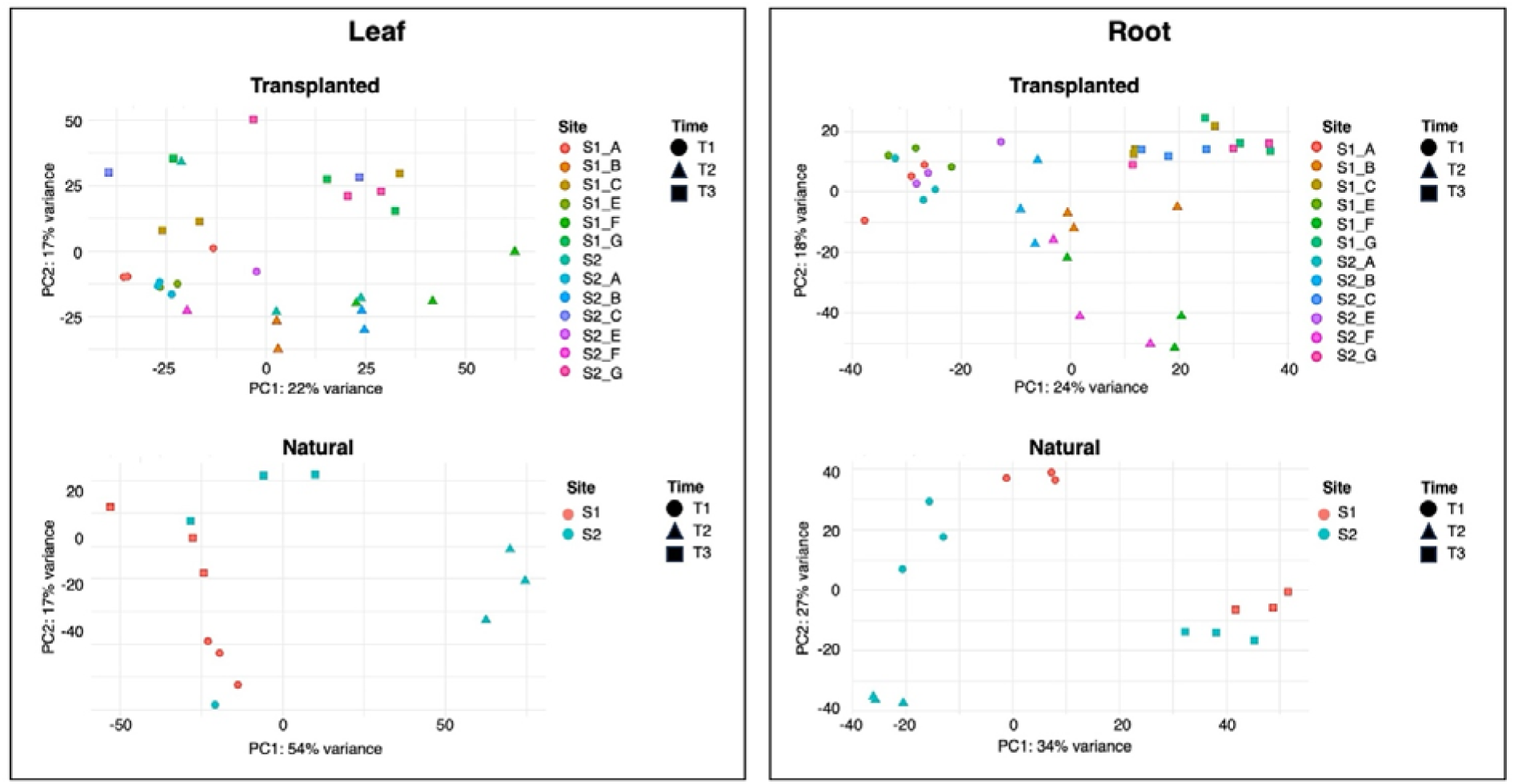
PCA of variance-stabilized RNA-seq data from root and leaf samples of transplanted and natural plants of *P. oceanica*. Samples are colored according to transplantation site and shaped according to time points (T1, T2, T3). The first two principal components are shown, with the percentage of explained variance reported on each axis. The absence of a consistent site-dependent clustering pattern supported pooling samples from the two transplantation sites in downstream differential expression analyses.

**Figure S7.**
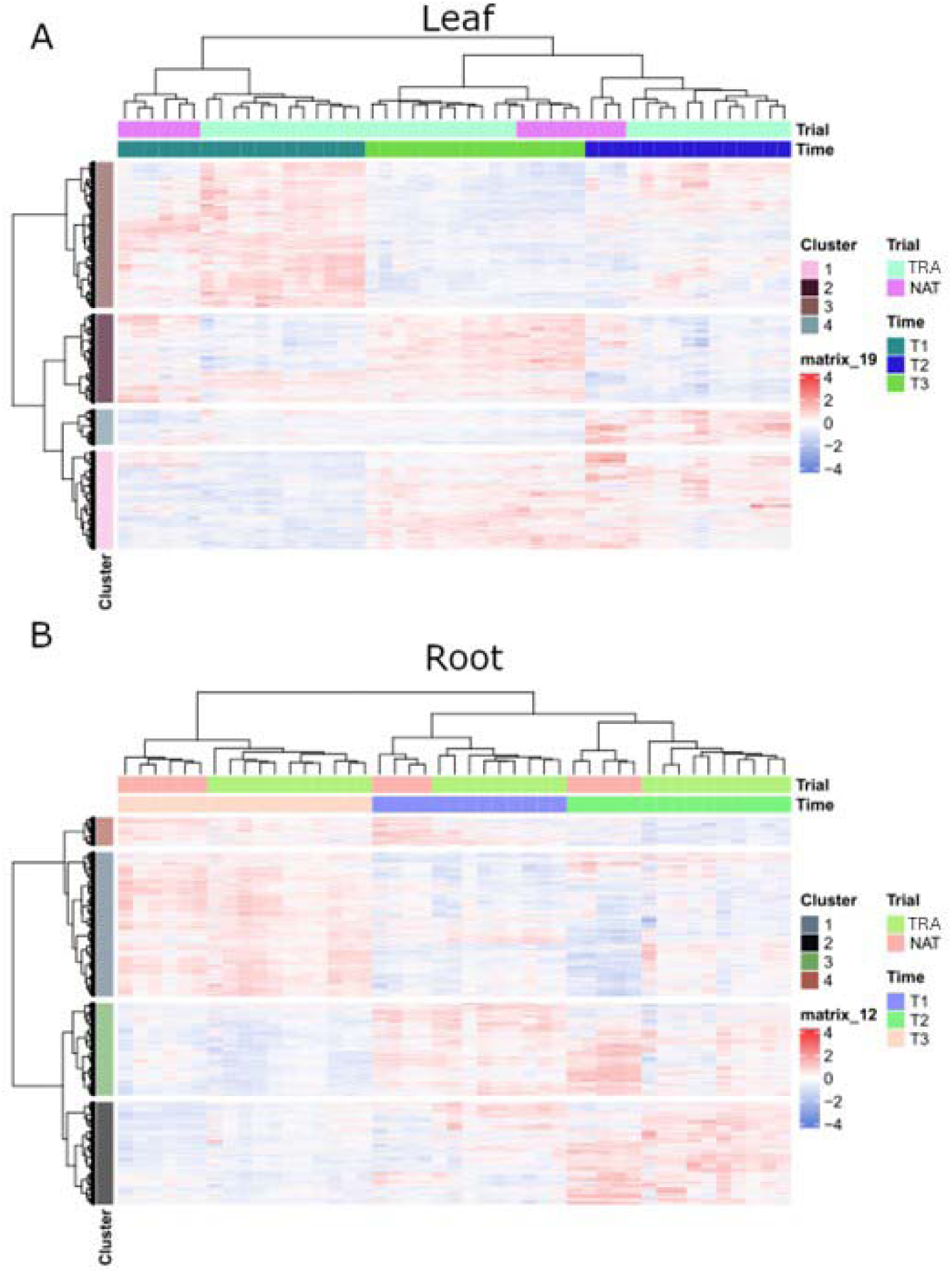
Hierarchical clustering and heatmap of DEGs. **A** Heatmap of the 6207 DEGs identified in leaf tissue. Genes are grouped into four clusters (1–4) based on their expression trends across time points (T1–T3) and plant types (TRA, NAT). **B** Heatmap of the 4564 DEGs identified in root tissue, partitioned into four clusters. For both panels, the top dendrograms and color bars indicate the clustering of samples by plant type and time point, while the color scale represents the z-score of scaled expression levels.

**Figure S8.**
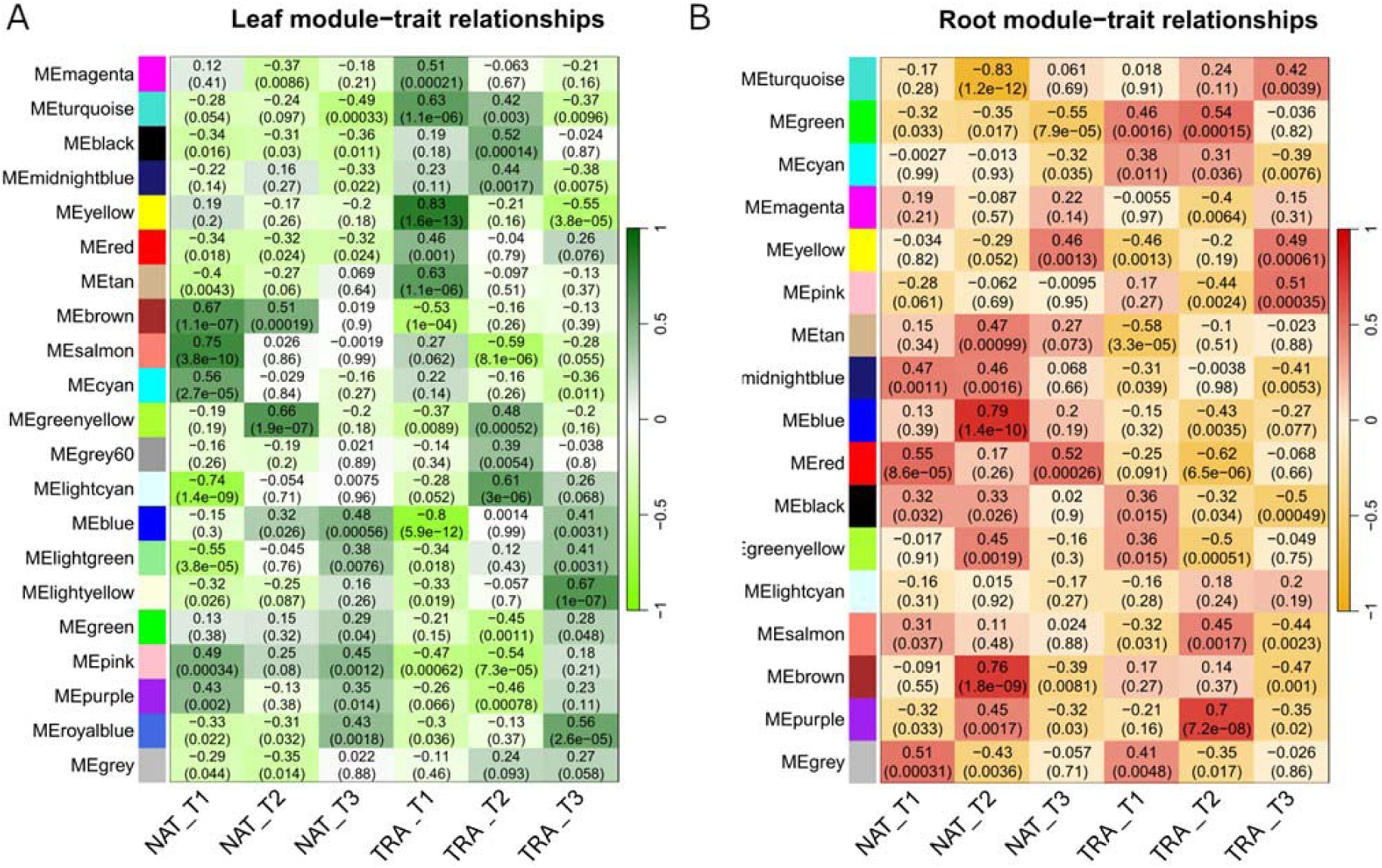
Complete module-trait association matrices for leaf and root datasets. Pearson correlation analysis between all co-expression modules and experimental traits (Time and Plant type) in leaf (**A**) and in root (**B**) samples. In both panels, the color scale indicates the correlation coefficient (green/red for positive and negative correlations), with the corresponding P-values reported in parentheses for each module-trait pair.

**Supplementary Table S1.** List of samples from transplanted (TRA) and natural (NAT) plants.

**Supplementary Table S2.** Water chemical and microbiological parameters at transplantation (T0) and collection times (T1-T3).

**Supplementary Table S3**. LRT analysis of differentially expressed genes. Summary of DEGs identified in leaf tissues across different time points and plant types.

**Supplementary Table S4**. LRT analysis of differentially expressed genes. Summary of DEGs identified in root tissues across different time points and plant types.

**Supplementary Table S5**. Hub genes identified within the selected co-expression modules. Detailed list of the top-connected genes (hubs) for the tissue-specific modules selected for the analysis.

**Supplementary Video S1**. Transplantation of seedlings in situ.

